# Auranofin shows bactericidal activity in *Pseudomonas aeruginosa* by targeting thiol homeostasis

**DOI:** 10.64898/2026.09.11.750932

**Authors:** Alexandre Luscher, Yves Mattenberger, Léna Falconnet, Natacha Civic, Christian van Delden, Thilo Köhler

## Abstract

The intrinsic resistance of *Pseudomonas aeruginosa* to many antibiotics is driven by the combined action of the outer membrane permeability barrier and multidrug efflux pumps, limiting the discovery of compounds active against this pathogen. To facilitate identification of antibacterial molecules with intracellular targets, we constructed a hyper-permeable *P. aeruginosa* PA14 strain lacking the four major Mex efflux systems and expressing the FhuA-derived hyperpore. This strain exhibited markedly increased susceptibility to diverse antimicrobials. Screening of 2,400 predominantly FDA-approved compounds identified the antirheumatic drug auranofin as a potent inhibitor which displayed bactericidal activity against the hyperpermeable strain. Selection of resistant mutants identified gain-of-function mutations in the MexQ efflux pump, indicating that altered substrate specificity of the MexP-MexQ-OpmE efflux system can reduce auranofin susceptibility. To investigate its mode of action, we analyzed mutants defective in the thioredoxin and glutathione redox systems. Whereas disruption of the thioredoxin pathway had no effect on susceptibility, glutathione-deficient mutants were hypersusceptible to auranofin. Exogenous glutathione restored resistance, and intracellular thiol measurements demonstrated that auranofin depletes the cellular thiol pool, consistent with direct neutralization by glutathione. These findings support a model in which auranofin disrupts glutathione-dependent thiol homeostasis in *P. aeruginosa*. More broadly, our study demonstrates that overcoming permeability and efflux barriers is an effective strategy to reveal antibacterial activities of approved drugs against Gram-negative pathogens.

## INTRODUCTION

The low outer membrane permeability combined with efficient tripartite drug efflux pumps represent the major barrier to antimicrobials in Gram-negative bacteria. A particularly challenging group of opportunistic pathogens are the Gram-negative non-fermenters (*Pseudomonas aeruginosa*, *Acinetobacter baumannii*, *Stenotrophomonas maltophilia*, *Burkholderia cepacia* complex), which in general are endowed with higher intrinsic resistance levels than Gram-negative enterobacterial species (1–3). Furthermore, their propensity to develop frequently multidrug resistance phenotypes has urged WHO to list *P. aeruginosa* and *A. baumannii* as priority organisms requiring urgent need for novel antibiotics or alternative treatment strategies (WHO) (4).

The outer membrane (OM) of Gram-negative bacteria is an asymmetric lipid layer composed of lipopolysaccharide (LPS) in the outer leaflet and phospholipids in the inner leaflet. In addition, surface charge distribution and LPS capping may modulate the OM permeability. Furthermore, the number and the structural architecture of porins for the facilitated diffusion of more hydrophilic compounds across the OM shows notable differences between Gram-negative bacteria. For instance in *P. aeruginosa* deletion of all non-essential porin genes has revealed that only the OM porin OprD is able to transport antibiotics, namely the carbapenems imipenem and meropenem (5)(6).

To overcome the low OM permeability, several strategies have been attempted including (i) OM permeabilization, (ii) efflux pump inhibition and (iii) high jacking of TonB-dependent transporters (TBDT) to actively transport siderophore-drug conjugates. This last strategy is exemplified by cefiderocol, a conjugate linking a siderophore catechol moiety to a modified ceftazidime antibiotic. This drug was approved for the treatment of MDR isolates of *P. aeruginosa* and *A. baumannii*. Cefiderocol circumvents the low OM permeability by flagging a β-lactam through vectorization for active uptake via siderophore transporters in Gram-negative bacteria (7–9). Further attempts to overcome the low OM permeability are the combination of Gram-positive drugs with membrane permeabilizers, as highlighted by conjugated substituted vancomycin derivatives(10, 11). However, these additions to the antibiotic armamentarium still belong to already existing drug classes (β-lactams, tetracyclines, glycopeptides). The screening efforts in the 90s of large antimicrobial compound libraries to identify novel antibiotic targets have revealed mainly the fatty acid and LPS synthesis inhibitors (12–14). A fundamental question thus remains unanswered: are there any novel drug targets in Gram-negative bacteria, but which have been missed so far in classical antimicrobial whole cell screenings, due to the particularly efficient permeability barrier of these critical Gram-negative pathogens.

To address this fundamental question, we have used here a hyperpermeable *P. aeruginosa* strain, by genetically removing the four clinically relevant drug efflux pump operons and by expressing a large ungated hyperpore in the OM (15, 16).

We screened two molecule libraries of FDA-approved drugs to screen for antibacterial activity in this hypersusceptible strain background. We identified auranofin as the most potent compound, which affects thiol homeostasis in *P. aeruginosa* pointing towards a novel mode of action and a novel backbone for medicinal chemistry modifications.

## RESULTS

### Antibiotic susceptibility of a hyper-permeable *P. aeruginosa* strain

We constructed a hyper-permeable derivative of the *P. aeruginosa* reference strain PA14, termed PA14-HP, by genetically deleting the loci for the clinically relevant Mex efflux pumps MexAB-OprM, MexCD-OprJ, MexXY, MexEF-OprN and by introducing a plasmid expressing the FhuA-hyperpore (FhuA^hyp^)(17). This hyperpore is derived from the ferrichrome transporter FhuA of *E. coli* (18), by genetically removing the plug domain and five extracellular loops thereby creating a beta barrel pore of 3 - 4 nm in diameter, which abrogates the outer membrane barrier providing access to the periplasmic space even for larger antimicrobial compounds (16, 17). FhuA^hyp^ is functional in several Gram-negative bacteria and has been used to assess the respective contributions of efflux pumps and outer membrane barrier to antibiotic susceptibility (15, 19)

Except for polymyxin-B, which targets the LPS of the OM, PA14-HP was hypersusceptible to all antimicrobials tested. Based on the MICs in the presence or the absence of efflux pumps, the hyperpore or both, all other tested antimicrobials could be classified into three categories, (i) molecules with high MW (>1000 dal) and active only on Gram-positive bacteria (vancomycin, bacitracin), (ii) molecules affected mainly by efflux (lincomycin, ciprofloxacin, sulfamethoxazole, trimethoprim) and (iii) those affected by both mechanisms (novobiocin, linezolid, tetracycline, ampicillin, aztreonam, azithromycin) (Table 1). Members of the last two groups had MW in the range of 290 to 749 dal. Multiplication of MIC ratios for individual contributions of either efflux or hyperpore resulted in similar ratios as those obtained for efflux and hyperpore vs wild-type. This suggests that both mechansims were additive rather than synergistic. Similar results were obtained in a PAO1 strain background (15). Rifampicin could not be evaluated since PA14 is intrinsically resistant to this antibiotic due to a mutation in the *rpoB* gene (20).

**Table 1.** Contribution of efflux pump deletion and/or hyperpore expression on antimicrobial susceptibility in *P. aeruginosa* PA14.

| Antibiotic | MIC (μg/ml) |  |  |  | fold change MIC<sup>2</sup> |  |  | MW (dal) |
| --- | --- | --- | --- | --- | --- | --- | --- | --- |
|  | PA14<br>pSRK-Gm | Δ4mex<br>pSRK-Gm | PA14<br>pFhuA-HP | Δ4mex<br>pFhuA-HP | efflux | hyperpore | efflux+<br>hyperpore |  |
| polymixin | 1 | 1 | 1 | 0.5 | **1** | **1** | **2** | 1386 |
| bacitracin | 1024 | 512 | 256 | 128 | **2** | 4 | 8 | 1422 |
| vancomycin | 1024 | 128 | 128 | 32 | 8 | 8 | 32 | 1450 |
| lincomycin | 1024 | 128 | 1024 | 128 | 8 | **1** | 8 | 406 |
| ciprofloxacin | 0.125 | 0.008 | 0.062 | 0.008 | 16 | **2** | 16 | 331 |
| sulfamethoxazole | 128 | 8 | 64 | 8 | 16 | **2** | 16 | 253 |
| trimethoprim | 128 | 2 | 64 | 1 | 64 | **2** | 128 | 290 |
| novobiocin | 256 | 16 | 64 | 8 | 16 | 4 | 32 | 612 |
| linezolid | 1024 | 32 | 128 | 32 | 32 | 8 | 32 | 337 |
| tetracyclin | 16 | 0.125 | 4 | 0.125 | 128 | 4 | 128 | 444 |
| ampicillin | 128 | 8 | 8 | 1 | 16 | 16 | 128 | 349 |
| aztreonam | 4 | 0.25 | 0.25 | 0.016 | 16 | 16 | 256 | 435 |
| azithromycin | 128 | 2 | 8 | 0.125 | 64 | 16 | 1024 | 749 |

<sup>1</sup> MIC ratios in bold indicate antibiotics which are not or only marginally affected by either efflux, hyperpore or both mechanisms; MW, molecular weight.
<sup>2</sup>Fold change MIC reflect the contribution of efflux (MICs PA14/MICs Δ4mex), hyperpore (MICs PA14/MICs PA14 pFhuA-HP) or both (MICs PA14/MICs Δ4mex pFhuA-HP)

### Screening of FDA approved compound libraries for antimicrobial activity

We used the PA14-HP strain to screen two compound libraries, available at our institution: the Prestwick (www.prestwickchemical.com) and the NINDS (National Institute of Neurological Disorders and Stroke, www.commondataelements.ninds.nih.gov) libraries. These include respectively 1280 and 1040 mainly FDA-approved drugs, as well as 80 molecules referenced as kinase inhibitors yielding a total of 2400 compounds. The screening was performed in technical duplicates in 384 well plates, using compounds at a final concentration of 100 μM. Growth of PA14-HP in the presence of these compounds was measured as optical density at 600 nm (OD_600_) after 24 h incubation at 37 °C in MHB medium. Out of the 2400 compounds, 234 molecules (9.75 %) classified as biocides (antibiotics, antiviral, antifungal, disinfectant) and 110 as non-biocide compounds (4.58 %) showed a >90% reduction in growth compared to the control conditions (Fig.1). Azithromycin (MIC of 128 μg/ml for the wild type strain PA14) was used as a control and showed 99% growth inhibition of the PA14-HP strain at 7.5 μg/ml (i.e. 10 μM) in all screening assays. For each plate, the Z’ value was in the range of 0.5 to 1.0, thereby validating the quality across all batch screenings.

**Fig. 1.**
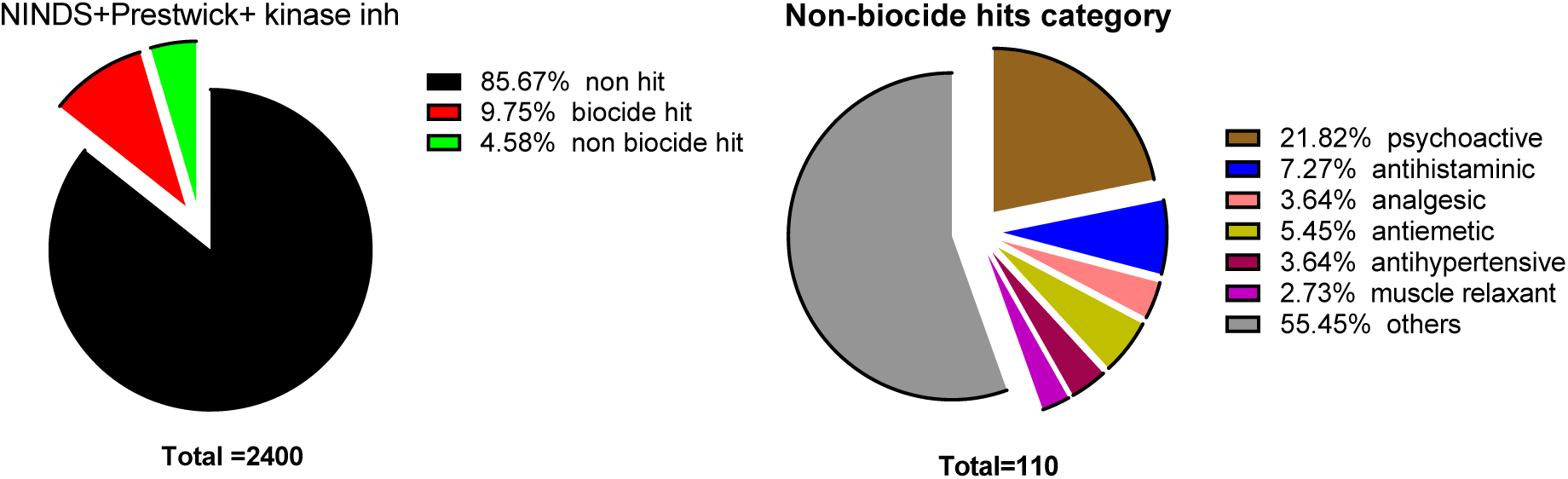
Screening of small compound drug libraries (NINDS, Prestwick, kinas inhibitors) in PA14-HP. The pie chart on the right is a blow-up of the non-biocide portion (green segment in left pie chart).

Twenty-nine of the 110 non-biocide compounds showed IC_50_ values below 20 μM and were considered as hits. Of these, twenty-one compounds showed at least 99% growth inhibition (Table S1). Seventeen of the 29 hits were retested under standard MIC conditions and nine showed MIC values ≤ 64 μM (Table S2). Three compounds, H-89 (N-[2-[[3-(4-Bromophenyl)-2-propenyl]amino]ethyl]-5-isoquinoline-sulfonamide dihydrochloride), clotrimazole and auranofin showed the highest activities with MICs of 4 μM, 4 μM and 8 μM, respectively. We discarded the sulphonamide derivative H-89 and the anti-fungal compound clotrimazole and focused our attention on auranofin, an FDA-approved drug for the treatment of rheumatoid arthritis (21).

### Auranofin activity is mainly affected by Mex efflux pumps in *P. aeruginosa*

We first determined whether auranofin was affected by drug efflux, OM permeability or both. The MIC values obtained in a set of isogenic PA14 mutants, showed that auranofin is a substrate of all four major Mex efflux pumps (MexAB-OprM, MexCD-OprJ, MexXY, MexEF-OprN), since deletion of individual pump operons or double or triple deletions did not increase the susceptibility of PA14 to auranofin (MIC of 256 μM). Only in the Δ4mex strain, auranofin MICs dropped to 16 μM (Table S3). Expression of the hyper-pore in the PA14 wild type or in the Δ4mex mutant did not further increase auranofin susceptibility, suggesting that efflux is mainly responsible for the intrinsic resistance to this compound. The data, however, clearly indicate that target(s) exist for this molecule in *P. aeruginosa*. To investigate whether auranofin was bacteriostatic or bactericidal, we performed killing assays, which showed a two-log reduction in viable counts of the Δ4mex strain after 24 h incubation in the presence of 128 x the MIC, compared to the vehicle control (DMSO). Hydrogen peroxide at 100 x the MIC showed a faster killing without detectable bacteria recovered after 6 h incubation (Fig. 2).

**Fig. 2.**
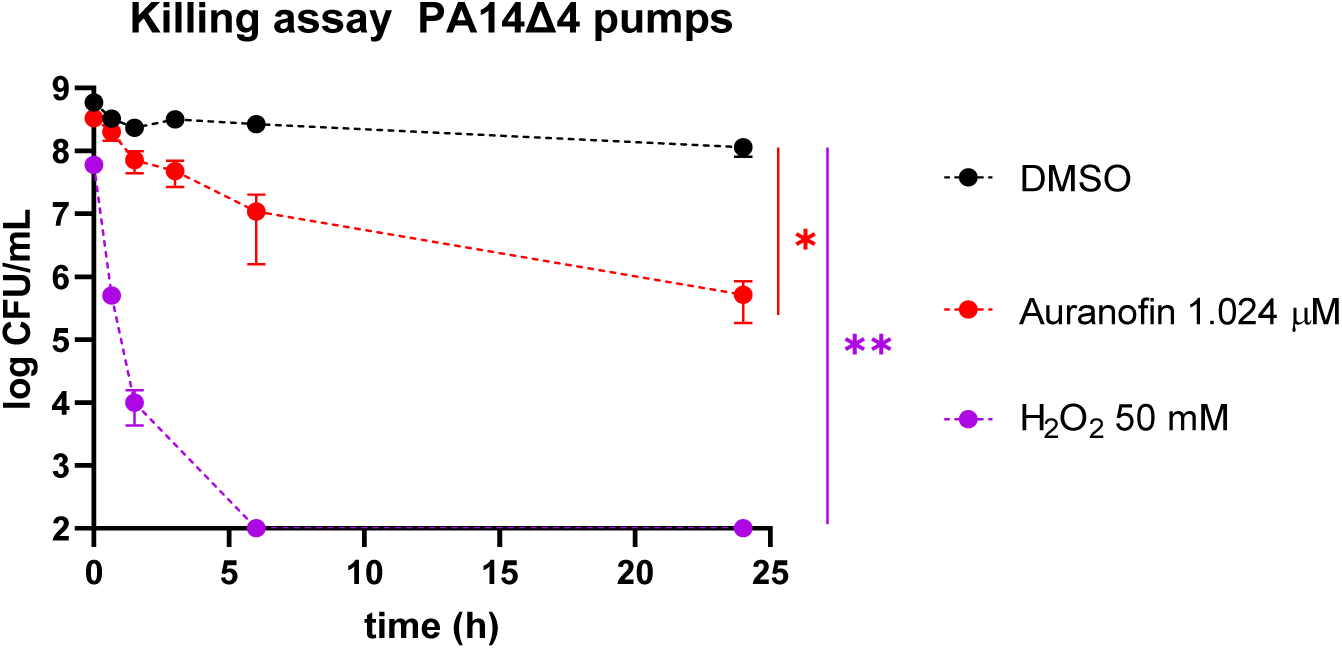
Auranofin is bactericidal in the efflux pump deficient strain. Killing assays were performed on stationary phase bacteria. Values represent mean and standard deviation of four independent experiments. Unpaired t-test, *, p<0.05, **, p<0.01

### Searching for auranofin target(s) in *P. aeruginosa*

To explore the mode of action and identify possible targets of auranofin, we sought to select auranofin-resistant mutants by exposing the PA14Δ4mex strain to increasing concentrations of this drug. After five rounds of selection, three mutants were obtained (A-R1, A-R2 and A-R3). These mutants maintained their resistance phenotype even after growth in the absence of selective pressure. The decrease in susceptibility was specific to auranofin, since MIC values of other antibiotics including aztreonam, azithromycin, ciprofloxacin and polymyxin-B were unaffected (Table 1).

Whole genome sequencing revealed a single nucleotide polymorphism (SNP) in each of the three mutants, which was located in the same gene annotated as *mexQ* (PA3522, PA14_18780) in the Pseudomonas genome database (pseudomonas.com) (22). The *mexQ* gene encodes the inner membrane efflux transporter of the MexP-MexQ-OpmE RND type efflux system (23). Mutant A-R1 harboured a nt change at codon 216 of *mexQ* (647A>T) resulting in a Gln to Leu substitution, while mutants A-R2 and A-R3 displayed a Val to Leu substitution at codon 1008 (3022G>C). To verify that the latter mutation was responsible for the observed decrease in susceptibility, we cloned the entire *mexPQ*-*opmE* operon from the PA14 wild-type and from mutant A-R2, carrying the mutated *mexQ* gene, into the expression vector pIApX2. Expression of the wild type operon *mexP*-*mexQ-opmE* in the Δ4mex strain did not significantly affect auranofin MICs or those of other antibiotics tested (Table 1), suggesting that overexpression alone was not sufficient to affect auranofin activity. However, overexpression of the mutated version harbouring the Val1008Leu substitution (MexQ*) increased specifically auranofin MICs by 4-fold compared to the vector control (Table 2). We surmise that the amino acid substitutions identified in the MexQ efflux pump protein variants are gain of function mutations, which alter the affinity of the efflux pump to accommodate the thio-sugar auranofin.

**Table 2.** Susceptibility of mutants selected in PA14Δ4mex upon auranofin exposure and genetic complementation.

| Strains | MIC (μg/ml) | | | | |
| --- | --- | --- | --- | --- | --- |
| | Auranofin (μM) | AZM | AZI | CIP | Pmx-B |
| Δ4mex | 16 | 0.25 | 2 | 0.06 | 1 |
| Δ4mex A-R1 | 64 | 0.5 | 4 | 0.03 | 1 |
| Δ4mex A-R2 | 128 | 0.5 | 2 | 0.03 | 1 |
| Δ4mex A-R3 | 128 | 0.5 | 2 | 0.03 | 1 |
| Δ4mex+plApX2 | 16 | 0.25 | 2 | 0.03 | 1 |
| Δ4mex+pmexPQ-opmE | 32 | 0.5 | 2 | 0.015 | 1 |
| Δ4mex+pmexPQ*-opmE | 128 | 0.5 | 4 | 0.03 | 1 |

AZM, aztreonam; AZI, azithromycin; CIP, ciprofloxacin; Pmx-B, polymyxin-B; \*mexQ indicates the mutated mexQ allele G3022C (V1008L)

### Auranofin perturbs thiol homeostasis in *P. aeruginosa*

The selection of mutants with decreased auranofin susceptibility indicated a possible resistance mechanism but did not allow identification of potential targets of auranofin. Based on previous data reported in *S. aureus* and *Mycobacterium tuberculosis*, we hypothesized that auranofin, due to its gold complexed thiol-group, might affect the intracellular redox homeostasis (24). In Gram-negative bacteria, intracellular thiols are provided mainly by the tripeptide glutathione (γ-L-glutamyl-L-cysteinyl-glycine) while thioredoxin, a protein of typically 12 kDa is the main thiol redox protein in Gram-positive bacteria (25). In *P. aeruginosa* glutathione synthesis occurs through the sequential action of glutamate-cysteine ligase (GshA, PA5203) and glutathione synthetase (GshB, PA0407) (26). Thioredoxin is encoded by *trxA* (PA5240), which is reduced by the cognate thioredoxin reductase encoded by *trxB1* (PA2616). An additional putative thioredoxin reductase orthologue is annotated as TrxB2 (PA0849), displaying 73% amino acid identity (86% similarity) with TrxB1 (pseudomonas.com) (22). To assess the possible involvement of the glutathione and the thioredoxin systems for the antibacterial activity of auranofin, we first deleted the *mexPQ*-*opmE* efflux operon in the Δ4mex strain, yielding strain PA14Δ5mex. In this background we then constructed mutants deficient in the thioredoxin and glutathione systems. Surprisingly, neither the PAO1 nor the PA14 transposon libraries contain a thioredoxin (*trxA*) mutant (27, 28), suggesting essentiality of thioredoxin in *P. aeruginosa*. Using a selectable marker (ΩSm cassette), we nevertheless succeeded in generating a *trxA*::ΩSm mutant in PA14 as well as in the Δ5mex background. We next tested the susceptibility of the generated mutants towards auranofin, hydrogen peroxide and the thiol-oxidizing reagent diamide (29). Neither the thioredoxin (TrxA, PA5240) nor the thioredoxin reductase (TrxB1, PA2616; TrxB2, PA0849) deletions affected auranofin or diamide MICs. In contrast, the *trxA* mutant was more susceptible to H_2_O_2_, while the *trxB1*/*trxB2* mutant showed increased H_2_O_2_ MICs (Table 3). This suggests that the thioredoxin system is neither required for auranofin activity (generation of H_2_O_2_ or other ROS requiring the thioredoxin system for inactivation), nor is it a likely target. In contrast, the thioredoxin system was required for H_2_O_2_ stress, since in the absence of thioredoxin susceptibility to H_2_O_2_ increased, while deletion of thioredoxin reductases (TrxB1/TrxB2), which restore free thiol groups, resulted in decreased H_2_O_2_ susceptibility (Table 3).

**Table 3.** Susceptibility of *P. aeruginosa* thioredoxin and glutathione mutants.

| Strains | MIC |  |  |
| --- | --- | --- | --- |
| | Auranofin<br>(μM) | H<sub>2</sub>O<sub>2</sub><br>(mM) | Diamide<br>(μg/ml) |
| PA14 | >256 | 0.5 | >512 |
| PA14*trxA*::ΩSm | 256 | **0.125** | ND |
| PA14Δ*trxB1*Δ*trxB2* | >256 | 1 | ND |
| PA14Δ*gshA* | **32** | 0.5 | 512 |
| PA14Δ*gshB* | **32** | 0.5 | 512 |
| PA14Δ*gshA*Δ*gshB* | **32** | 0.5 | 512 |
| Δ5mex | 16-32 | 0.25 | 512 |
| Δ5mex-*trxA*::ΩSm | 16 | **0.06** | ND |
| Δ5mexΔ*trxB1*Δ*trxB2* | 32 | **1** | 512 |
| Δ5mexΔ*gshA* | **4** | 0.25 | **128** |
| Δ5mexΔ*gshB* | 8 | 0.125 | 256 |
| Δ5mexΔ*gshA*Δ*gshB* | **4** | 0.125 | **128** |

Values in bold indicate ≥ or ≤ 4-fold changes in MIC compared to PA14 or Δ5mex strain

Surprisingly, we observed the inverse situation with the glutathione mutants (*gshA*, *gshB*), which showed 4 to 8-fold decreased auranofin MICs, but were unaffected in their susceptibility to H_2_O_2_. This was true not only in the Δ5mex background but also in the PA14 wild type, which now displayed the same auranofin MICs as the Δ5mex mutant. Similar results were obtained with diamide, which oxidizes two glutathione (GSH) molecules to form a GSSH dimer. The *gshA* and *gshB* mutants were again more susceptible in the Δ5mex background (Table 3). GshA deletion also increased susceptibility to the aminoglycosides gentamicin and tobramycin (Table S4). Auranofin and H_2_O_2_ susceptibilities for the deletion mutants could be reverted by trans complementation with the corresponding gene. As expected, *gshA* could not complement *gshB* and vice versa (Table S5). From these data, we conclude that glutathione rather than thioredoxin antagonizes the antibacterial activity of auranofin in *P. aeruginosa*.

Several hypotheses can explain these results. We envisage that auranofin binds directly to glutathione and serves as a decoy for this redox molecule. We therefore performed MICs in the presence and absence of externally supplied glutathione. Indeed, 2 mM glutathione drastically increased MICs of auranofin in the Δ*gshA*Δ*gshB* double mutant, whether in the PA14 or in the Δ5mex strain background. Susceptibilities to aztreonam, included as a control, were not affected by glutathione (Table 4).

**Table 4.** Susceptibility of glutathione mutants to auranofin and aztreonam in presence of external glutathione.

| Strains | MIC Auranofin (μM) | | MIC AZM (μg/ml) | |
| --- | --- | --- | --- | --- |
|  | no GSH | 2mM GSH | no GSH | 2mM GSH |
| PA14 | >128 | >256 | 8 | 8 |
| $\Delta gshA\Delta gshB$ | 32 | >256 | 4 | 4 |
| $\Delta 5mex$ | 32 | >256 | 0.25 | 0.25 |
| $\Delta 5mex\Delta gshA\Delta gshB$ | 8 | 256 | 0.25 | 0.25 |

GSH, glutathione; AZM, aztreonam

We next measured the intracellular thiol concentration using the thiol reactive compound 5,5′-dithiobis-(2-nitrobenzoic acid) (DTNB, Ellman’s reagent). Addition of sub-MIC auranofin concentrations (32 μM) to PA14 wild type during growth reduced intracellular thiol concentrations by 2 to 3-fold, corresponding to the level measured in the glutathione deficient mutant Δ*gshA*Δ*gshB* (Fig. 3). In contrast, supplementing the growth medium with 2 mM glutathione increased intracellular thiol concentrations by 2-fold. The increase in intracellular thiol concentrations could be counteracted by simultaneously providing auranofin (32 μM), suggesting a direct interaction between these two compounds (Fig. 4, Fig. S1).

**Fig. 3.**
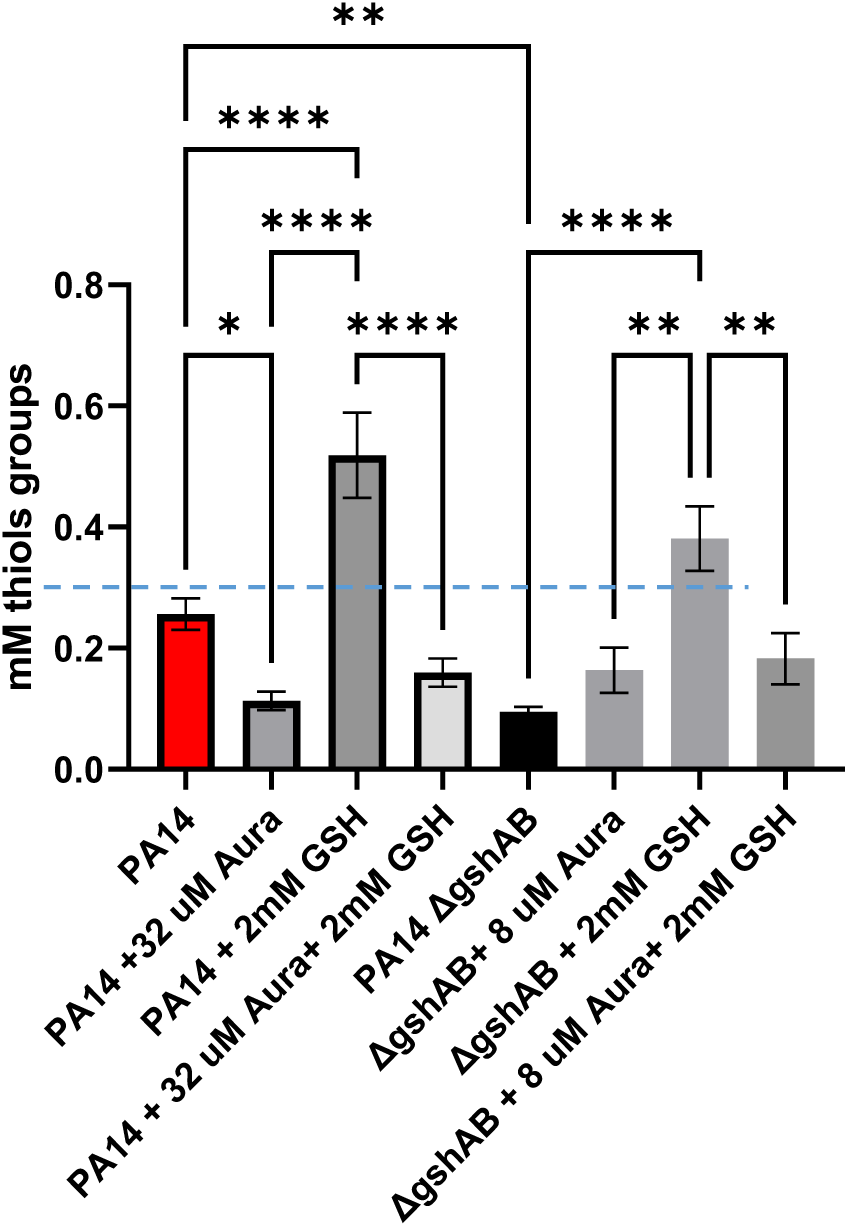
Measurement of intracellular thiol concentrations in PA14 and glutathione deficient mutant (Δ*gshAB*) grown in the presence or absence of auranofin (Aura) and/or glutathione (GSH) at sub-inhibitory concentrations. Stippled line indicates the detection limit of the assay. Statistical analysis was performed with the student t-test (Graph Pad Prism, v8.0). Values represent the average and standard deviations of three biological replicates. p-values: *, 0,05; **, 0.01; ***, 0.005; ****, 0.001.

**Fig. 4.**
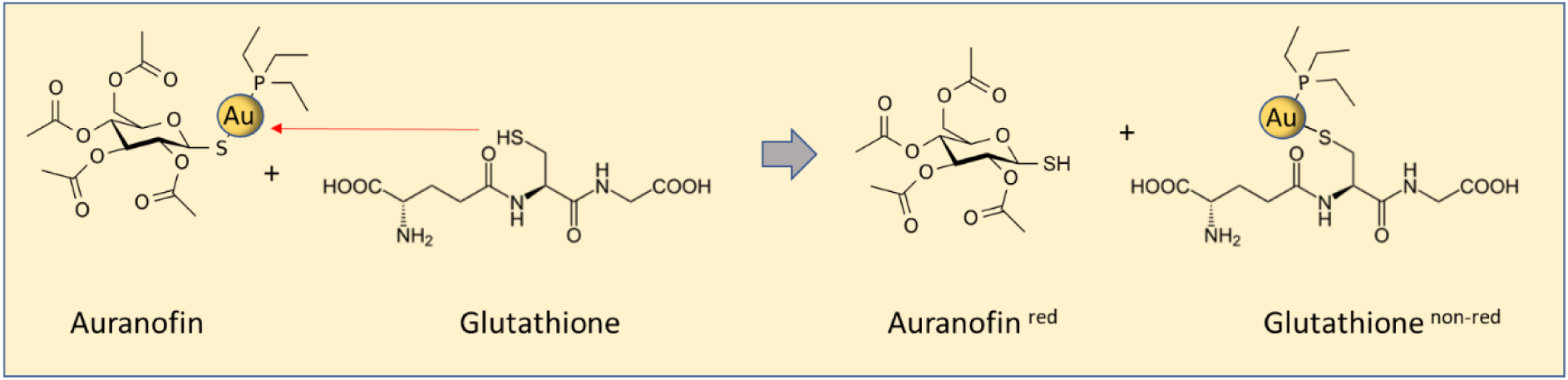
Proposed direct interaction between auranofin and glutathione.

Similar results were obtained in the glutathione deficient mutant. Growth in the presence of extracellular glutathione also increased intracellular thiol concentrations, which were reduced again by supplementing simultaneously auranofin at sub-MIC concentrations (Fig. 3). The detection limit for intracellular thiols in our assay was approximately 0.1 mM, corresponding to the levels of the Δ*gshA*Δ*gshB* mutant. This suggests that glutathione is the dominant thiol-providing compound in *P. aeruginosa*.

In summary our data suggest a direct interaction of auranofin with cytosolic glutathione, but not thioredoxin, resulting in disruption of thiol redox homeostasis (Fig. 4).

## DISCUSSION

In this study, we addressed the fundamental question of whether Gram-negative bacteria with intrinsically low outer membrane (OM) permeability harbor unexplored antimicrobial targets that remain inaccessible to conventional drug discovery approaches. To maximize intracellular access of candidate compounds, we constructed a hyperpermeable *Pseudomonas aeruginosa* strain lacking the four major Mex multidrug efflux systems and expressing a heterologous OM hyperpore. As expected, this strain exhibited markedly increased susceptibility to all antibacterial agents tested, including the Gram-positive-specific antibiotics vancomycin and bacitracin. These large peptide antibiotics (MW around 1,500 dal) were affected mainly by the OM barrier, whereas susceptibility to most other antibiotics was primarily determined by Mex-mediated efflux or by the combined action of efflux and the OM barrier. These findings are consistent with previous studies demonstrating the complementary roles of efflux pumps and the OM in the intrinsic resistance of *P. aeruginosa* (15, 19).

We exploited this hypersusceptible PA14-HP strain to screen two libraries comprising approximately 2,400 FDA-approved or clinically used compounds, including several established antibiotics that served as internal controls. In this hypersusceptible strain background, 1.2% of non-biocide molecules showed >90% growth inhibition compared to the vehicle control with IC_50_ values below 20 μM. This hit rate substantially exceeds those reported from large-scale whole-cell screens of *Mycobacterium abscessus* (10,000 compounds; hit rate 0.01 %) (30), *P. aeruginosa* PAO1 (230,000 compounds, hit rate 0.002%) (31) and *S. aureus* (250,000 compounds, hit rate 0.1 %) (32). Although our set of molecules was limited to FDA-approved or off-label drugs and partially biased by the presence of known antibiotics, our hit rate was almost 1,000-fold higher than the one reported for the whole cell screening with the *P. aeruginosa* wild type strain PAO1 (31). This fact clearly supports our initial hypothesis that potential drug targets are present in *P. aeruginosa* but remain inaccessible due to the permeability barrier imposed by the combination of efflux pumps and the OM.

Among the compounds identified, auranofin emerged as the most potent hit. Auranofin is an FDA-approved gold-containing drug originally developed for the treatment of rheumatoid arthritis (21), and has subsequently been shown to possess potent antibacterial activity against several Gram-positive pathogens, including *Staphylococcus aureus* and *Mycobacterium tuberculosis* (24). In contrast, auranofin showed only weak activity against *K. pneumoniae, A. baumannii* and *P. aeruginosa* (24). Our results demonstrate that the poor activity of auranofin against *P. aeruginosa* is not due to the absence of a susceptible cellular target but instead reflects intrinsic resistance mediated by the combined action of the outer membrane permeability barrier and multiple Mex efflux systems. Interestingly, the contribution of the OM barrier was apparent only in the wild-type background, whereas disruption of efflux pumps largely abolished this effect. These findings suggest that intracellular accumulation of auranofin is primarily limited by active efflux and that reduced OM permeability becomes restrictive only when efflux systems remain functional.

Selection of mutants with reduced susceptibility to auranofin identified mutations in the inner membrane transporter MexQ of the MexPQ-OpmE efflux system (23). One substitution (Q216L) is located within the large periplasmic domain of MexQ, whereas the second substitution (V1008L) affects the first residue at the periplasmic surface of the final transmembrane helix. Notably, this valine residue is highly conserved among several RND transporters, including MexB, MexD, MexF, and MexY, suggesting an important structural or functional role. The observation that only the MexPQ_V1008L_-OpmE variant pump but not the wild-type pump, conferred increased resistance suggests that auranofin is unlikely to be an efficiently transported physiological substrate of MexPQ-OpmE. Instead, the V1008L substitution may alter substrate specificity or transport efficiency, thereby enabling enhanced export of auranofin. Nevertheless, auranofin induced expression of the *mexPQ-opmE* operon in the Δ4mex strain, suggesting that the compound either shares structural features with endogenous MexPQ substrates or triggers the accumulation of metabolites that activate the MexPQ efflux pump. Further work will be required to distinguish between these possibilities and to clarify the molecular basis of MexPQ induction.

Although our mutant selection strategy did not identify the direct molecular target of auranofin, our data support previous observations that the compound disrupts thiol-dependent redox homeostasis (24). Indeed, auranofin altered intracellular thiol levels in a manner consistent with perturbation of the glutathione and thioredoxin redox systems. The increased intracellular thiol content observed upon addition of auranofin in the Δ*gshA*Δ*gshB* mutant may reflect compensatory upregulation of the thioredoxin system in response to the absence of glutathione. An inverse correlation was observed in A2780 cancer cells, where auranofin-induced depletion of thioredoxin resulted in increased glutathione production (33). These observations suggest overlapping functions of these two main thiol redox systems in Gram-negative bacteria. Because glutathione contributes not only to oxidative stress resistance but also to the regulation of pyoverdine and pyocyanin production as well as swimming and twitching motility in *P. aeruginosa* (26), interference with glutathione homeostasis may have consequences beyond bacterial viability. It is therefore conceivable that, at subinhibitory concentrations, auranofin may additionally function as an anti-virulence agent by attenuating multiple glutathione-dependent pathogenic traits.

Overall, our study demonstrates that overcoming the intrinsic permeability barrier of *P. aeruginosa* substantially expands the accessible antibacterial target space and enables the identification of compounds that would otherwise be overlooked in conventional whole-cell screens. The identification of auranofin as a potent inhibitor of the hypersusceptible strain provides proof of principle that clinically approved compounds can possess effective antibacterial activities against *P. aeruginosa* once permeability barriers are bypassed. These findings further emphasize that improving compound uptake and retention may be as important as identifying new molecular targets in the development of future therapies against multidrug-resistant Gram-negative pathogens.

## ACKNOWLEDGMENTS

This study was funded by CONFIRM grant No. RC04-06 from the HUG Private Foundation of the Geneva University Hospitals to T.K, C.v.D and P. Viollier (University of Geneva, Switzerland). We also acknowledge funding from the BRDIGE project No. 40B2-0_211759. We thank Y. Cambet (READs platform of the Faculty of Medicine, University of Geneva) for performing the initial library screenings. We also thank Prof. L. Movileanu (Syracuse University, USA) for sharing the *fhuAh* sequence for subcloning purposes.

## AUTHOR CONTRIBUTIONS

Conceptualization and methodology, A.L., T.K., N.C. and C.v.D.; Investigation A.L., Y.M., L.F., N.C. and T.K.; Writing, A.L., T.K. and C.v.D.; Funding Acquisition, T.K. and C.v.D.; Supervision, T.K. and C.v.D.

## SUPPLEMENTAL INFORMATION

Supplemental information can be found online at:

### Data and code availability

Code and raw data supporting the conclusions of this study can be found at zenodo.org; DOI: 10.5281/zenodo.22081640

## MATERIAL AND METHODS

### Bacterial growth conditions

Bacteria strains were grown in Lysogeny Broth at 37°C with agitation (250 rpm) or as otherwise stated. Strains and plasmids used in this study are listed in Table S6 and primers in Table S7. Antibiotic stock solutions were prepared in double-distilled H_2_O or in Dimethyl sulfoxide (DMSO). Auranofin was purchased from SIGMA and dissolved in DMSO (stock solution: 34 mg/ml).

### Screening of molecule libraries

NINDS and Prestwick compounds collections were provided by the READS platform (University of Geneva, Switzerland). Test compounds were screened at 100 μM final concentration (from DMSO 100 mM stock solutions) in 384-well plates containing 40 μl of MHB broth supplemented with 2.5 μg/ml gentamycin and 2mM IPTG for plasmid maintenance and hyperpore expression, respectively. Approximately 10^4±0.5^ bacteria were added to each well, and plates were incubated at 37°C for 24h without shaking. Azithromycin at 10 μM was used as inhibition control and DMSO (4 μl/well) as vehicle control to allow Z-factor calculation for each plate. After incubation, optical density at 600 nm (OD_600_) was measured with a Synergy Biotek H1 plate reader.

### Minimal inhibitory concentration (MIC) determinations

MICs were determined by broth double microdilution method in Müller-Hinton Broth (MHB) (Becton Dickinson, Franklin Lakes, NJ, USA), in accordance with the Clinical and Laboratory Standards Institute (CLSI) guidelines. If not indicated otherwise, glutathione (SIGMA) was added to MHB medium at 2mM final concentration.

### Selection of mutants with decreased auranofin susceptibility

To select auranofin-resistant mutants in the PA14Δ4mex strain background, serial passages with increasing concentrations of auranofin were performed in MHB in technical triplicates. After MIC determination, a 50 μl aliquot of the last well showing visible growth (0.5 x MIC) was regrown in 2 ml of LB for 8 h, before being challenged with serial dilutions of auranofin. After six passages an 8-fold increase in auranofin MICs was achieved. A five μL aliquot of the last well showing visible growth from each triplicate experiment was spread on LB agar plates. Individual colonies were picked, regrown without selection and tested by MIC determinations to confirm stable and clonal resistance to auranofin.

### Construction of knockout mutants

The construction of unmarked deletion mutants was performed as described by Hoang et al (34). DNA fragments of 550-650 bp length corresponding to upstream and downstream regions of the gene to be deleted were PCR-amplified using primers F1/R1 & F2/R2 (Table S6), and cloned into the suicide vector pEXG2(35). PCR amplification was achieved with Q5 high fidelity DNA polymerase (NEB) under the following conditions: 98°C for 2 min, then 30 cycles of 98°C for 20s, 63°C for 30s, 72°c for 30s and a final extenstion for 4 min at 72°C. *E. coli* strain DH10B was used as a host for all cloning experiments. Plasmid constructs were introduced in target strains of *P. aeruginosa* by biparental mating using *E. coli* strain ST18. The following steps of mutant generation were carried out as previously described (36). The generated chromosomal deletions were verified by PCR amplification with external primers F1/R2 and subsequent Sanger sequencing of the generated amplicons.

### Construction of expression plasmids for genetic complementation

For complementation of the genetic deletion mutants, the corresponding coding region including at least 50 bp upstream of the start codon were PCR amplified from genomic DNA using Q5 high fidelity DNA polymerase (NEB) under the following conditions: 98°C for 2 min, then 30 cycles of 98°C for 20s, 63°C for 30s, 72°c for 30s/kb, and a final extension for 4 min at 72°C. Amplicons were cloned into the expression vector pIApX2 using restriction enzymes *Bam*HI and *Hin*dIII (NEB). Newly-constructed vectors were then inserted into *P. aeruginosa* strains by electroporation and transformants were selected on LB plates supplemeneted with carbenicilin at 200 μg/ml (wt strains) or 50 μg/ml (Δmex strains). Constructs were verified by Sanger sequencing.

### Measurement of free-thiol groups

The intracellular concentration of free thiol groups was measured using Ellman’s reagent: 5,5-dithio-bis-(2-nitrobenzoic acid) (DTNB) (Merck-SIGMA, Switzerland). Strains were grown for 18 h in LB medium as described above. A 1 ml culture sample was centrifuged at 9’000 rpm for 5 min, the supernatant was discarded and the pellet resuspended in 0.5 ml of 0.01M HCl. Suspensions were quickly vortexed, left 10 min at RT and centrifuged again at 9’000 rpm for 5 min. In a 96-well plate (TPP 92096), 50 μl of the collected supernatant were added to 50 μl of test solution (110 mM Na_2_HPO_4_, 40 mM NaH_2_PO_4_, 15 mM EDTA, 0.3 mM DTNB), and finally 50 μl of stop solution (50mM Imidazole, 15mM HCl, 1mM EDTA). Free thiol groups were colorimetrically assessed by measuring the optical density at 405 nm, using a Biotek Synergy H1 plate reader. A standard curve of reduced glutathione (Merck-SIGMA) was prepared as 10-fold serial dilutions (0.005 to 5 mM) and treated together with the samples.

### Killing assay

Stationary-phase grown bacteria were exposed to auranofin or hydrogen peroxide in 1 ml NaCl 0.9%. Viable cell counts were determined at different timepoints by spotting serial dilutions (in NaCl 0.9% supplemented with 50 mM sodium thiosulfate) on LB agar plates.

### Whole Genome Sequencing

Genomic DNA was isolated using Qiagen DNeasy kit according to the manufacturer’s instructions, from a 2 mL LB medium grown for 16-18h (OD_600_ ≈ 2). Whole genome sequencing was performed at the iGE3 Genomic Platform of the University of Geneva. Libraries were constructed using the Nextera protocol followed by 50 bp paired-end sequencing on an Illumina HiSeq instrument. DNA sequencing reads were mapped on the UCBPP-PA14 genome (acUKH-supercont1.1). The genomic variants were called with FreeBayes v1.1.0. The variant annotation and effect prediction was performed with SnpEff v.4.3. Mutations identified by variant call were confirmed after PCR amplification of the corresponding DNA region and Sanger Sequencing of the amplicon.

### Statistical analyses

All statistical analysis was performed with GraphPad Prism version 8.0. Experiments were performed in three biological replicates with at least two technical duplicates. Statistical significance is reported in the figure legends, and data are presented as mean ± SD as indicated.

