## Supplementary Tables and Figures for "Auranofin shows bactericidal activity in *Pseudomonas aeruginosa* by targeting thiol homeostasis"

### Supplementary material

**Table S1:** Excel file showing IC<sub>50</sub> values for compounds showing > 95% growth inhibition on PA14Δ4mex

See raw data in zenodo repository at DOI: [10.5281/zenodo.22081640](https://doi.org/10.5281/zenodo.22081640)

**Table S2.** MIC values of PA14-HP for compounds with IC<sub>50</sub> < 20 μM

| Compounds | MIC (μM) |
| --- | --- |
| Cytidine | >100 |
| Puromycin | 12.5 |
| Tannic acid | >100 |
| Vitamin B3 | >100 |
| 8-OH quinoline | 50 |
| Kynurenic acid | >256 |
| Phenazopyridine·HCl | 128 |
| Clomipramine·HCl | 64 |
| α-cyano-4-OH-cinnamic acid | >256 |
| Ciclopirox | 32 |
| Clotrimazole | 4 |
| α-cyano-3-OH-cinnamic acid | >512 |
| H-89* | 4 |
| Zn-pyrrhione | 32 |
| Phenazopyridine | 64 |
| Sildenafil | >512 |
| **Auranofin** | **8** |

\* H-89, N-[2-[[3-(4-Bromophenyl)-2-propenyl]amino]ethyl]-5-isoquinolinesulfonamide

**Table S3.** Susceptibility of Mex efflux pump mutants towards auranofin

| Strains | MIC Auranofin (μM) |
| --- | --- |
| PA14 | >256 |
| PA14 + pSRK-fhuAh | 64 |
| ΔmexAB-oprM | 256 |
| ΔmexXY | >256 |
| ΔmexCD-oprJ | >256 |
| ΔmexEF-oprN | >256 |
| ΔmexAB-oprM ΔmexXY | 256 |
| ΔmexAB-oprM ΔmexCD-oprJ | 256 |
| ΔmexXY ΔmexCD-oprJ | >256 |
| ΔmexAB-oprM ΔmexXY ΔmexCD-oprJ | 256 |
| ΔmexAB-oprM ΔmexXY ΔmexCD-oprJ ΔmexEF-oprN | 16 |
| ΔmexAB-oprM ΔmexXY ΔmexCD-oprJ ΔmexEF-oprN<br>+ pSRK-fhuAh | 4-8 |

**Table S4.** Susceptibility of glutathione mutants to antibiotics

| PA14-derived strains | MIC (mg/L) |  |  |  |  |  |  |
| --- | --- | --- | --- | --- | --- | --- | --- |
|  | GEN | TET | CIP | AMP | CAZ | AZT | PMB |
| PA14 | 1 | 16 | 1 | >256 | 2 | 128 | 1 |
| ΔgshA | 0.25 | 16 | 1 | >256 | 2 | 128 | 1 |
| ΔgshB | 0.5 | 16 | 2 | >256 | 4 | 128 | 1 |
| ΔgshAΔgshB | 0.25 | 16 | 1 | >256 | 2 | 64 | 1 |
| Δ5mex | 0.5 | 0.125 | 0.06 | 16 | 1 | 2 | 1 |
| Δ5mexΔgshA | 0.125 | 0.125 | 0.125 | 32 | 2 | 2 | 1 |
| Δ5mexΔgshB | 0.25 | 0.125 | 0.125 | 32 | 1 | 2 | 1 |
| Δ5mexΔgshA<br>ΔgshB | 0.125 | 0.125 | 0.125 | 32 | 1 | 2 | 1 |

GEN, gentamicin; TET, tetracyclin; CIP, ciprofloxacin; AMP, ampicillin; CAZ, ceftazidime; AZT, azithromycin; PMB, polymyxin B

**Table S5.** Susceptibility of thioredoxin and glutathione deficient mutants

| Strain | MICs |  |
| --- | --- | --- |
|  | Auranofin<br>(μM) | H<sub>2</sub>O<sub>2</sub><br>(mM) |
| PA14 | 256 | 0.5 |
| *trxA::*ΩSm + pSRKGm | 256 | 0.125 |
| *trxA::*ΩSm + ptrxA | 256 | 0.5 |
| Δ*gshA* + pIApX2 | 32 | 0.5 |
| Δ*gshA* + pgshA | >256 | 0.5 |
| Δ*gshA* + pgshB | 32 | 0.5 |
| Δ*gshB* + pIApX2 | 32 | 0.5 |
| Δ*gshB* + pgshA | 64 | 0.5 |
| Δ*gshB* + pgshB | >256 | 0.5 |
| Δ*gshA* Δ*gshB* + pIApX2 | 32 | 0.5 |
| Δ*gshA* Δ*gshB* + pgshA | 32 | 0.5 |
| Δ*gshA* Δ*gshB* + pgshB | 64 | 0.5 |
| Δ*gshA* Δ*gshB* + ptrxA | 32 | 0.5 |
| Δ*gshA* Δ*gshB* + ptrxB1 | 32 | 0.5 |
| Δ*gshA* Δ*gshB* + ptrxB2 | 32 | 0.5 |

**Table S6.** Strains and plasmids used in this study

| Strains/plasmids | Relevant characteristics | Reference, source |
| --- | --- | --- |
| ***P. aeruginosa*** |  |  |
| PA14 | Reference strain | (20) |
| $\Delta mexAB-oprM$ | unmarked deletion of *mexAB-oprM* | D. Pletzer |
| $\Delta gshA$ | unmarked deletion of *gshA* | This study |
| $\Delta gshB$ | unmarked deletion of *gshB* | This study |
| $\Delta gshAgshB$ | unmarked deletions of *gshA* and *gshB* | This study |
| PA14 |  |  |
| $\Delta 4mex$ | unmarked deletions of *mexAB-oprM mexCD-oprJ mexEF-oprN mexXY* | This study |
| $\Delta 4mex-AR.1$ | $\Delta 4mex$ Auranofin-resistant selected clone, *mexQ*<sup>A647T</sup> | This study |
| $\Delta 4mex-AR.2$ | $\Delta 4mex$ Auranofin-resistant selected clone, *mexQ*<sup>G3022C</sup> | This study |
| $\Delta 4mex-AR.3$ | $\Delta 4mex$ Auranofin-resistant selected clone, *mexQ*<sup>G3022C</sup> | This study |
| $\Delta 5mex$ | $\Delta 4mex$ + unmarked deletion of *mexPQ-opmE* | This study |
| $\Delta 5mex\ trxA::\Omega Sm$ | $\Delta 5mex$ + Sm<sup>R</sup>-marked deletion of *trxA* | This study |
| $\Delta 5mex\ \Delta trxB1-B2$ | $\Delta 5mex$ + unmarked deletions of *trxB1*, *trxB2* | This study |
| $\Delta 5mex\ trxA::\Omega Sm\ \Delta trxB1-B2$ | $\Delta 5mex\ trxA::\Omega Sm$, unmarked deletions of *trxB1*, *trxB2* | This study |
| $\Delta 5mex\ \Delta gshAgshB$ | $\Delta 5mex$ + unmarked deletions of *gshA*, *gshB* | This study |

### ***E. coli***

|  |  |  |
| --- | --- | --- |
| ST18 | *pro thi hsdR*<sup>+</sup> Tmp<sup>r</sup> Sm<sup>r</sup>; chromosome::RP4-2<br>Tc::Mu-Kan::Tn7/λpirΔ*hemA* | (37) |

### **Plasmids**

|  |  |  |
| --- | --- | --- |
| pEXG2 | gene replacement vector, Gm-R | (35) |
| pSRKGm | IPTG-inducible broad-host range expression<br>vector, Gm-R | (38) |
| pfhuAh | fhuAh (hyperpore) sequence with a C-term 6xHis tag (fhuA-ΔC/Δ5L-6xHis)<br>cloned into pSRKGm | (39) |
| pIApX2 | Broad-host range expression vector, Ap-R | I. Attree (Grenoble,<br>France) |
| pHP45ΩSm | Source of Sm<sup>R</sup> cassette, SmR, Ap-R | (40) |
| pgshA | *gshA* from PA14 cloned into pIApX2, Ap-R | This study |
| pgshB | *gshB* from PA14 cloned into pIApX2, Ap-R | This study |
| pmexPQ-opmE | *mexPQ-opmE* from PA14 cloned into pIApX2,<br>Ap-R | This study |
| pmexPQ*-opmE | *mexPQ*<sup>G3022C</sup>-*opmE* from Δ4*mex*AuraR.2 cloned<br>into pIApX2, Ap-R | This study |
| pEXG2-Δ*mexCD*-<br>oprJ | 1.2-kb fusion fragment of the up- and<br>downstream region of *mexCD-oprJ* from PA14,<br>Gm-R | This study |
| pEXG2-Δ*mexEF*-<br>oprN | 1.2-kb fusion fragment of the up- and<br>downstream region of *mexEF-oprN* from PA14,<br>Gm-R | This study |
| pEXG2-Δ*mexPQ*-<br>opmE | 1.2-kb fusion fragment of the up- and<br>downstream region of *mexPQ-opmE* from PA14,<br>Gm-R | This study |
| pEXG2-Δ*mexXY* | 1.2-kb fusion fragment of the up- and<br>downstream region of *mexXY* from PA14, Gm-R | This study |
| pEXG2-Δ*gshA* | 1.2-kb fusion fragment of the up- and<br>downstream region of *gshA* from PA14, Gm-R | This study |
| pEXG2-ΔgshB | 1.2-kb fusion fragment of the up- and downstream region of *gshB* from PA14, Gm-R | This study |
| pEXG2-ΔtrxB1 | 1.2-kb fusion fragment of the up- and downstream region of *trxB1* from PA14, Gm-R | This study |
| pEXG2-ΔtrxB2 | 1.2-kb fusion fragment of the up- and downstream region of *trxB2* from PA14, Gm-R | This study |
| pEXG2-trxA::ΩSm | 2.2-kb Sm<sup>R</sup>-cassette cloned between the up- and downstream region of *trxA* from PA14 1.2-kb fusion fragment, Gm-R, Sm-R | This study |

**TABLE S7.** Primers used in this study

| Primer | Sequence (5' - 3') | Purpose |
| --- | --- | --- |
| mexXY-hind-F1 | ACACAAGCTTGCACATCGCCAGACAGACCT | Deletion *mexXY* |
| mexXY-xba-R1 | ACACTCTAGAACGTCCTGGCCTTCCTCGTA | Deletion *mexXY* |
| mexXY-xba-F2 | ACACTCTAGAGAACGCCATCCTCATCATCG | Deletion *mexXY* |
| mexXY-xho-R2 | ACACCTCGAGGATCCGCTCGGTAGCCTGAC | Deletion *mexXY* |
| mexCDJ-hind-F1 | ACACAAGCTTGTCCGGGCGGTACTGGAATA | Deletion *mexCD-oprJ* |
| mexCDJ-bam-R1 | ACACGGATCCGAACTCAGCGCCAGGGACTC | Deletion *mexCD-oprJ* |
| mexCDJ-bam-F2 | ACACGGATCCTCGACAACCACCTGCGCTAC | Deletion *mexCD-oprJ* |
| mexCDJ-eco-R2 | ACACGAATTCGTACCCTCGAACGCCTCACC | Deletion *mexCD-oprJ* |
| mexEFN-hind-F1 | ACACAAGCTTCAAGCGCAAGGTGGTCCTG | Deletion *mexEF-oprN* |
| mexEFN-bam-R1 | ACACGGATCCCGGTGAATTCGTCCCACTC | Deletion *mexEF-oprN* |
| mexEFN-bam-F2 | ACACGGATCCGAAGGCACCACCGATTTCCT | Deletion *mexEF-oprN* |
| mexEFN-eco-R2 | ACACGAATTCCCCACCAACAGACCAACAG | Deletion *mexEF-oprN* |
| mexPQE-F1-hind | ACACAAGCTTCCTCGAGCTTCACGGTCAAA | Deletion *mexPQ-opmE* |
| mexPQE-R1-bam | ACACGGATCCTTCTCTACCAGGCGGCCTTC | Deletion *mexPQ-opmE* |
| mexPQE-F2-bam | ACACGGATCCCTGCAGGAGACCGATGATGC | Deletion *mexPQ-opmE* |
| mexPQE-R2-eco | ACACGAATTCAACGCAGAGGCACAGGAGTG | Deletion *mexPQ-opmE* |
| mexPQE-F-bam | ACACGGATCCTTGCCGGACTTCCCTTCCTA | Complementation<br>*mexPQ-opmE* |
| mexPQE-R-hind | ACACAAGCTTAACGCAGAGGCACAGGAGTG | Complementation<br>*mexPQ-opmE* |
| gshA-hind-F1 | ACACAAGCTTCGCTGTTCTCGCTGATGGAC | Deletion *gshA* |
| gshA-bam-R1 | ACACGGATCCGTGGGCGTGATGAACTCCAG | Deletion *gshA* |
| gshA-bam-F2 | ACACGGATCCATGCGGAACTGACGCCTTC | Deletion *gshA* |
| gshA-eco-R2 | ACACGAATTCCTGGAAGAGCAGGGCATCCT | Deletion *gshA* |
| gshA-bam-F | ACACGGATCCGCTCGGTCCACCCTCATATTG | Complementation *gshA* |
| gshA-hind-R | ACACAAGCTTAGCAGCGTAGAGCGGCTACC | Complementation *gshA* |
| gshB-hind-F1 | ACACAAGCTTGGACTTGAACGCGCTGTTGT | Deletion *gshB* |
| gshB-bam-R1 | ACACGGATCCCGAGCGAGCTGTCCTTCTTG | Deletion *gshB* |
| gshB-bam-F2 | ACACGGATCCGAAATCAAGGACGGCGACAA | Deletion *gshB* |
| gshB-xba-R2 | ACACTCTAGACCAGGGTCTTGCTGATCTGG | Deletion *gshB* |
| gshB-bam-F | ACACGGATCCCCATAATGCCCGGAACAGG | Complementation *gshB* |
| gshB-hind-R | ACACAAGCTTGGGGGCGGAAAAGTCTATGA | Complementation *gshB* |
| trxB1-hind-F1 | ACACAAGCTTCCGAGCGTCATGAACAGGAT | Deletion *trxB1* |
| trxB1-bam-R1 | ACACGGATCCTTGTCGACTTCGGTGGTGGT | Deletion *trxB1* |
| trxB1-bam-F2 | ACACGGATCCCGAACACCGACCTGTTCCAG | Deletion *trxB1* |
| trxB1-xba-R2 | ACACTCTAGACTGGTAGGCCAGTTGCATCG | Deletion *trxB1* |
| trxB2-hind-F1 | ACACAAGCTTGCGCCTACGGTATCGAGCA | Deletion *trxB2* |
| trxB2-bam-R1 | ACACGGATCCCTGCATCCGCTGCATCAGT | Deletion *trxB2* |
| trxB2-bam-F2 | ACACGGATCCTGAAGGACGGCTACCTGGTG | Deletion *trxB2* |
| trxB2-eco-R2 | ACACGAATTCGCAGGTTCTCCACGCAGTTG | Deletion *trxB2* |
| trxA-hind-F1 | ACACAAGCTTTAGATCAGGCGGGCTTCCTC | Deletion *trxA* |
| trxA-bam-R1 | ACACGGATCCTGCTCGAAGCTGGCATCAGT | Deletion *trxA* |
| trxA-bam-F2 | ACACGGATCCGCCTTCCTCGACGCCAATA | Deletion *trxA* |
| trxA-eco-R2 | ACACGAATTCCGAAGTAGCGTTCGCCCTCT | Deletion *trxA* |
| mexQ-R | ACTGCTCTTCGTCGCTCAGG | Verify *mexQ* mutation |
| mexQ-F | GATGACCTCGCTGGCGTTC | Verify *mexQ* mutation |

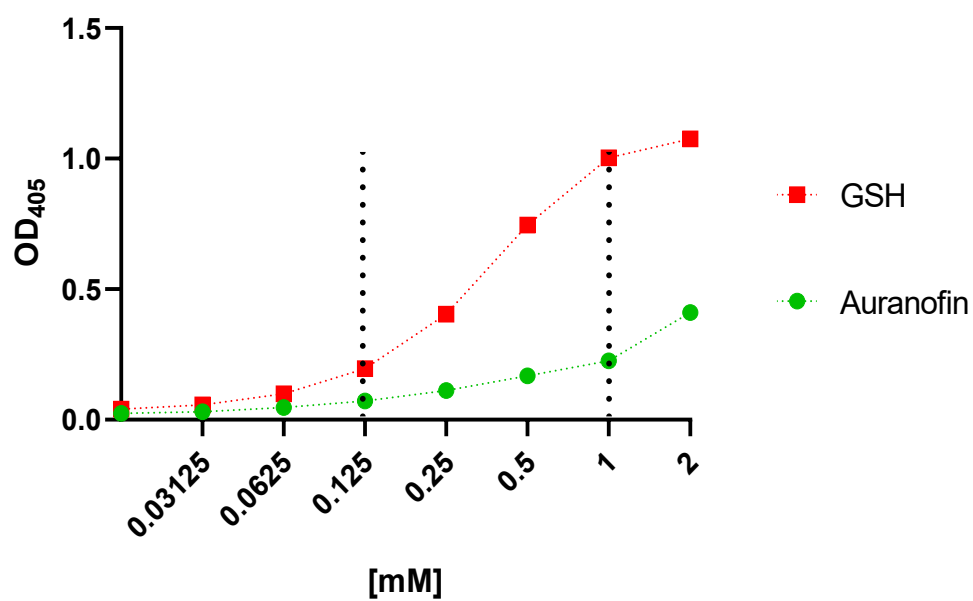

**Fig. S1.** Reaction of DTNB with auranofin and glutathione. Vertical dotted lines delineate the linear range for auranofin or glutathione concentrations used to measure free thiols in samples shown in Fig. 4. Auranofin was added at 32 μM to PA14 and at 8 μM to PA14Δ4mex (Fig. 4). At these concentrations the absorption contributed by auranofin does not affect measures of free intracellular thiols.
